# Grace-AKO: A Novel and Stable Knockoff Filter for Variable Selection Incorporating Gene Network Structures

**DOI:** 10.1101/2022.03.30.486361

**Authors:** Peixin Tian, Yiqian Hu, Zhonghua Liu, Yan Dora Zhang

## Abstract

**Motivation:** Variable selection is a common statistical approach to identifying genes associated with clinical outcomes of scientific interest. There are thousands of genes in genomic studies, while only a limited number of individual samples are available. Therefore, it is important to develop a method to identify genes associated with outcomes of interest that can control finite-sample false discovery rate (FDR) in high-dimensional data settings.

**Results:** This article proposes a novel method named Grace-AKO for graph-constrained estimation (Grace), which incorporates aggregation of multiple knockoffs (AKO) with the network-constrained penalty. Grace-AKO can control FDR in finite-sample settings and improve model stability simultaneously. Simulation studies show that Grace-AKO has better performance in finite-sample FDR control than the original Grace model. We apply Grace-AKO to the prostate cancer data in The Cancer Genome Atlas (TCGA) program by incorporating prostate-specific antigen (PSA) pathways in the Kyoto Encyclopedia of Genes and Genomes (KEGG) as the prior information. Grace-AKO finally identifies 47 candidate genes associated with PSA level, and more than 75% of the detected genes can be validated.

**Availability and implementation:** We developed an R package for Grace-AKO available at: https://github.com/mxxptian/GraceAKO

## 1 Introduction

Identifying genes and pathways associated with complex traits is a primary focus for advancing the scientific understanding in genomic studies, for which extensive clinical experiments and genetic counseling are required (Katsevich and Sabatti, 2019). A major challenge is that genomic data is usually high-dimensional but with a limited sample size. Currently, there are rich public genomic data, such as the Kyoto Encyclopedia of Genes and Genomes (KEGG, https://www.genome.jp/kegg/), which provides gene-regulatory pathways including biological regulatory relationships between genes or gene products. These gene-regulatory pathways form a network that can be represented as a graphical structure, where the vertices are genes or gene products, and the edges are gene-regulatory pathways. Inspired by the nature of genomic data, graph-constrained estimation (Grace) offers a novel perspective on variable selection for genomic data by taking graphical structures into account. The predictors in the network are gene-expression data linked by gene-regulatory pathways for a specific clinical outcome, which can also be called graph-structured covariates (Li and Li, 2010). These graphical structures, such as gene-regulatory pathways, can improve the sensitivity of detecting pathways (Rahnenführer et al., 2004). Therefore, Li and Li (2008) developed a Grace model, network-constrained regularization procedure by encoding the graphical structures into a Laplacian matrix to incorporate this kind of prior biological information into regression analysis. The network-constrained regularization procedure includes the Lasso (Tibshirani, 1996), and the elastic net regulation procedure (Zou and Hastie, 2005) as special cases. Particularly, the network-constrained penalty may be transformed to a Lasso-type problem by constructing an augment dataset, which can also retain the automatic variables selection property (Li and Li, 2008). The augment dataset can extend the sample size from *n* to *n* + *p*, allowing the model to choose *p* variables despite the fact that *n* ≪ *p*. Moreover, because the loss function of the network-constrained penalty is a convex function, it can ensure the grouping effect of the regression in the case of identical predictors. In accordance with the theorem of Li and Li (2008), the quantitative description of the grouping effect is measured beyond a half of the elastic net model 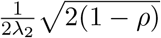, where *λ*_2_ is a fixed scalar and *ρ* is the correlation between the vertices, which means that if two vertices are highly correlated (e.g., *ρ* = 1), the difference between their coefficient paths could be almost 0. According to Katsevich and Sabatti (2019), the optimal policy for this variable selection effort would be to identify significant relevant genes and provide error control for these discoveries (both genes and variants). However, these conventional regression approaches only control false discovery rate (FDR) asymptotically with no guarantee in finite-sample settings. Specifically, there is a lack of effective approaches for variable selection which can not only integrate the graphical structure but also provide a guarantee of finite-sample FDR control.

To address this challenge, we propose Grace-AKO, a novel method for identifying genes associated with complex traits of scientific interest that integrates the core concept of aggregation of multiple knockoffs (AKO, Nguyen et al. (2020)) with the network-constrained regularization procedure (Li and Li, 2008). The salient idea of knockoff inference is to generate knockoff variables by mimicking the correlation structure of the original variables without considering the response variable (conditionally on the original variables) (Candes et al., 2018). Model-X knockoffs, as opposed to fixed-X knockoffs, regarded the original variables as random and relied on the specific stochastic properties of the linear model, thus extending knockoff inference to high-dimensional data (Candes et al., 2018). These knockoff variables are applied to control finite-sample FDR served as negative controls so that the original variables are selected if they are considerably more connected with the response variable than their knockoff variables. Specifically, the knockoff inference uses various types of feature statistics to determine which variables are significant and which are not. The feature statistics impose a flip-sign property, which implies that swapping the variables and their knockoffs alters the sign of the feature statistics. The methods for constructing the feature statistics are i.i.d. random for the “null hypothesis” whose coefficients are zero (Barber and Candès, 2015a; Candes et al., 2018). To control FDR, Barber and Candès (2015a) developed a data-dependent threshold whose derivation formula may be regarded as an estimate of the fraction of false discoveries. In addition, variables whose feature statistics are larger than the threshold may be selected, and estimations of the FDR can be converted to provide finite-sample FDR control with a high degree of accuracy. However, model-X knockoffs were generated using Monte Carlo sampling, which made it challenging to reproduce the results. Thus, multiple knockoffs were proposed to address this limitation, allowing for a more stable finite-sample FDR control and more reproducible findings (Emery and Keich, 2019; Gimenez and Zou, 2019; Nguyen et al., 2020). In particular, statistical aggregation is a typical statistical approach to solve instability by aggregating model-X knockoffs inference. Aggregation of multiple knockoffs (AKO) was proposed by Nguyen et al. (2020), which rested on a reformulation of model-X knockoffs to introduce an intermediate feature statistic. It brought the idea from Meinshausen et al. (2009) to replicate model-X knockoff procedure multiple times, and then performed statistical aggregation to generate new intermediate feature statistics. Hence, it is more stable than model-X knockoff filter.

Specifically, the key contribution of our proposed Grace-AKO is that we integrate the graphical structure to conduct variable selection with finite-sample FDR control. Based on the knockoff inference property, we update the Laplacian matrix in the network-constrained penalty and perform variable selection with the original explanatory variables and their knockoffs. The primary steps of Grace-AKO are summarized as follows. First, we generate model-X knockoffs according to the correlation structure of the graph-structured covariates (Candes et al., 2018) and encode the graphical structures into a Laplacian matrix (Chung, 1997).Second, we simultaneously fit the graph-structured covariates and their model-X knockoffs into the network-constrained regularization procedure to multiple feature statistics: Lasso coefficient-differences (LCDs). Third, we repeat the above procedures multiple times and employ the statistical aggregation approach (Meinshausen et al., 2009) to transform the LCDs into new intermediate feature statistics, Aggregated Grace Coefficients (AGCs). Fourth, we conduct the Benjamini-Hochberg (BH) procedure (Benjamini and Hochberg, 1995) on the multiple AGCs to select the candidate variables. In our simulation studies, we show that Grace-AKO has satisfactory performance, allowing for higher reproducibility of results, and can control the FDR in finite-sample settings. We further analyze a prostate cancer data set from The Cancer Genome Atlas (TCGA) program using Grace-AKO, and then identify 47 candidate genes, of which 75% were also found in the previous literature.

The remainder of this article is organized as follows. In Section 2, we describe the method of Grace-AKO under the linear regression framework. In Section 3, we assess the performance of Grace-AKO using simulation studies. In Section 4, we apply Grace-AKO to a prostate cancer data set from the TCGA program by incorporating the KEGG pathways as prior information. In Section 5, we briefly summarize our method.

## 2 Method

In genomic studies, we usually apply a regression model to identify genes and pathways associated with the trait of interest by linking high-dimensional data (e.g., microarray gene-expression data) to the trait. Consider the following linear model where ***X*** is a *n* × *p* design matrix with *n* observations and *p* predictors, and **y** is the response:

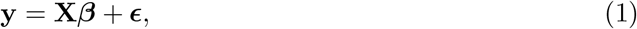

where the design matrix **X** = (**x**_1_,…, ***x***_p_) and **x**_j_ = (x_1*j*_, x_2*j*_,…, *x*_n*j*_), *j* = 1,…,*p* represents a vector of the graph-structured covariates from genomic data (e.g., gene-expression data), and the coefficient ***β*** represents the contribution of the graph-structured covariates to the trait of interest, and **ϵ** is a vector of random errors. We further assume that the response is centred and the predictors are standardized,

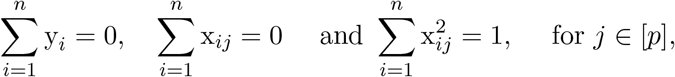

where [*p*] denotes the set including {1, 2,…, *p*}. The main goal is to estimate ***β*** = (*β*_0_, *β*_1_,…, *β_p_*) and select the predictors with nonzero contribution. The regulatory relationships of biological networks for some complex traits, represented as graphical structures between genes or gene products in genomic studies, shed light on underlying biological knowledge, where the covariates are the graph’s nodes and the edges indicate functional relationships between two genes. The biological networks can be utilized to identify the differentially expressed genes (Wei and Li, 2007, 2008). Specifically, in such a graphical structure, the genes are linked by edges with certain probabilities, where the edge probability is interpreted as a weight in an undirected graph to form a weighted graph (Li and Li, 2010). To incorporate the prior information about the biological networks,Li and Li (2008) proposed a network-constrained regularization criterion:

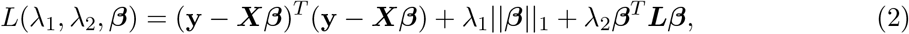

where ***L*** is a non-negative Laplacian matrix of a weighted graph containing biological networks information, and || · ||_1_ indicates the *L_1_* norm, and *λ*_1_ is the Lasso penalty (Tibshirani, 1996), and *λ*_2_ is the penalty for the Laplacian matrix ***L***. Specifically, the Laplacian matrix was first introduced by Chung (1997), which included numerous properties of the graph by its consistent set of the eigenvalues or spectrum. When *p* is large, the model (1) is treated as “sparse”, in which most elements of the coefficient ***β*** are zero (Li and Li, 2010). In equation (2), the L_1_ norm deals with sparse matrices, and ***β*^T^*Lβ*** induces a smooth solution of coefficients of the graph-structured covariates. Additionally, ***L*** depicts the graphical structure assuming that set *V* includes vertices corresponding to the graph-structured covariates, and *W* is the weights of the edges in which *w*(*u*, *v*) denotes a weight of the edge between the graph-structured covariates *u* and *v*, and the degree of vertex *v* is represented as *d_v_* = ∑_*u*~*v*_ *w*(*u*, *v*). In the genomic data, *w*(*u*, *v*) quantifies the uncertainty of the edge between two vertices, such as the probability of an edge connecting two graph-structured covariates when the graphical structure is constructed from data. Motivated by Chung (1997), we apply the normalized ***L*** (Li and Li, 2008):

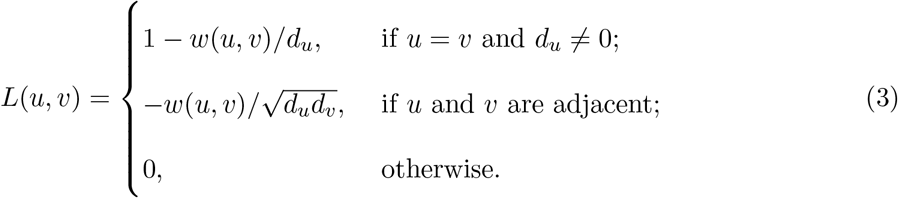

Therefore, we can rewrite the second penalty term ***β**^T^**Lβ*** of equation (2) as follows (Li and Li, 2008):

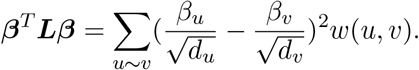

The network-constrained regularization procedure integrates the known biological network’s information for variable selection. By introducing the network-constrained penalty, the network-constrained regularization procedure was able to identify more interpretable genes and subnetworks related with the outcome of interest, while simultaneously inducing sparsity and smoothness of the biological network and the coefficients. However it does not ensure false dis-covery rate (FDR) control in finite-sample settings (Li and Li, 2008). The network-constrained regularization procedure only has the asymptotic property only when *n* → ∞ and *p* is fixed and suffers from identifying numerous false positive discoveries when *p* is large and the number of samples *n* is limited.To address this issue, we introduce knockoff inference and multiple knockoffs to control finite-sample FDR to achieve a stable performance. The knockoff filter procedure was first proposed in Barber and Candès (2015b), and Candes et al. (2018) further proposed model-X knockoff filter to extend its application to high-dimensional data. Model-X knockoffs, 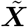 are generated from the original data by Monte Carlo sampling and retain the same data structure as the originals ***X***, in which ***X*** and 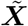 are pairwise exchangeable. We summarize the properties of model-X knockoffs as in Candes et al. (2018):

1. Swapping the locations of related elements, **x**_*j*_ and 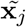, would not change the joint distribution of 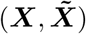 conditional on **y**.
2. Once the original covariates **X** are known, their model-X knockoffs, 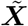 provides no extra information on the response variable **y**.

The knockoffs filter is a cheap method to control finite-sample FDR since it does not require strong assumptions about the design matrix **X**. Due of the random nature of model-X knockoffs sampling, however, the outcome would be unstable and cannot be guaranteed to be reproduced. To increase the stability, multiple knockoffs approaches were developed, and aggregation of multiple knockoffs (AKO) is one of them (Nguyen et al., 2020). In this article, we present a novel method for variable selection termed Grace-AKO, which combines the biological network information for improved variable selection with finite-sample FDR control using knockoff filter technique. Our proposed Grace-AKO has four major steps.

First, we generate model-X knockoffs, 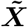 from the original data matrix **X** using R package “knockoff” (Candes et al., 2018). 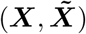 is further regarded as a new design matrix of the predictors. Based on the properties of model-X knockoffs whose correlation structure is the same as that of the original variables, we generate a new normalized Laplacian matrix ***L***. In summary, we generate knockoff variables and update the Laplacian matrix to include the graphical structure of knockoff variables in this step. The knockoff variables are introduced into the model based on their properties.

Second, the response variable ***y*** and the new predictors 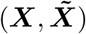 are fitted in equation (2).

Inspired by Li and Li (2008), a natural solution of Grace-AKO is equivalent to the following optimization problem:

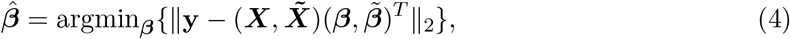

subject to:

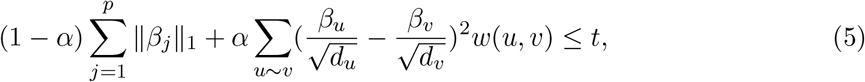

where *α* = *λ*_2_/(*λ*_1_+*λ*_2_), and *t* is a constant value, and 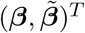 is a vector of the new coefficients. Furthermore, in the function 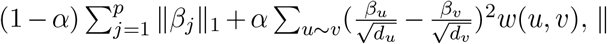, || · ||_1_ deals with sparse data matrix consistent with its function in the network-constrained penalty, and the second term penalizes the weighted sum of the squares of the difference of coefficients between the graph-structured covariates, which is scaled by the degree of the associated vertices in the network. Specifically, when two genes are connected, it is expected that their coefficients would be similar rather than identical, which is accomplished by applying the second term of the penalty (Li and Li, 2008).

To identify relevant variables, we compute the Lasso coefficient-difference (LCD) as the feature statistic to measure the evidence against null hypothesis (*β_j_* = 0) (Candes et al., 2018):

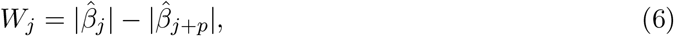

where 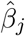, and 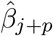 are the estimated coefficients of **x**_j_ and 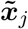, respectively. Due to the symmetric distribution of *W_j_* under the null, *W_j_* equally takes on positive and negative values (Candes et al., 2018). Moreover, a large positive value of *W**j*** suggests that the distribution of **y** is statistically dependent on **x**_*j*_ and that there is a strong probability that **x**_*j*_ is a relevant gene associated with the response **y**. In this step, the network-constrained penalty is integrated with knockoff variables to control finite-sample FDR. The natural solution of Grace-AKO follows the same optimization problem as the network-constrained regularization procedure. However, the model-X knockoffs procedure only generates knockoff variables once by Monte Carlo sampling, which leads to instability. To solve this challenge, we repeat the knockoff generation process multiple times and apply the statistical aggregation strategy to increase stability (Meinshausen et al., 2009).

Third, following Nguyen et al. (2020), Grace-AKO transforms the feature statistic *W_j_* into a new intermediate feature statistic *q_j_*:

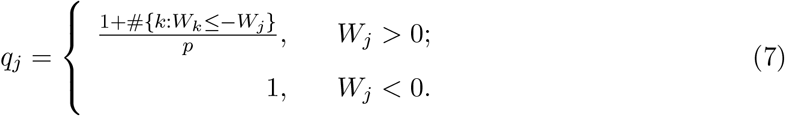

We repeat the aforementioned steps *B* times, including the generation of knockoffs and the calculation of the intermediate feature statistic *q_j_*, to generate a *B* × *p* matrix of the intermediate feature statistics. Then, we propose a new feature statistic, Aggregated Grace Coefficient (AGC), which is derived by applying the quantile aggregation algorithm (Meinshausen et al., 2009) to the *B* × *p* matrix:

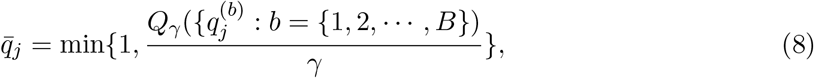

where *γ* is the quantile point, and *Q*(·) denotes the quantile function, and *B* is the pre-specified replication times. To summarize the third step, we generate model-X knockoffs *B* times and then use the quantile aggregation procedure to yield AGCs, 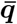. The quantile aggregation approach introduces intermediate feature statistics and aggregated feature statistics, which are based on the concept of statistical aggregation (Meinshausen et al., 2009) to improve stability.

Fourth, we apply the Benjamini-Hochberg procedure (BH) (Benjamini and Hochberg, 1995) to the AGCs, 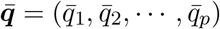 to compute a data-dependent threshold:

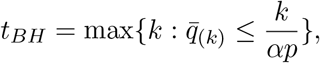

where *α* is the user-specified nominal FDR level. We finally choose the candidate variables satisfying the following requirement:

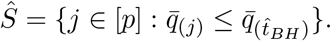

In this article, we measure the performance using the modified false discovery rate (mFDR):

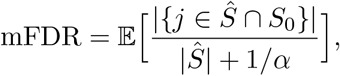

where *S*_0_ = {*j* ∈ [*p*]: *β_j_* = 0} includes the predictors that have no effect on the trait of interest.

The implementation details are available in Algorithm 1.

### Algorithm 1: Algorithm of Grace-AKO

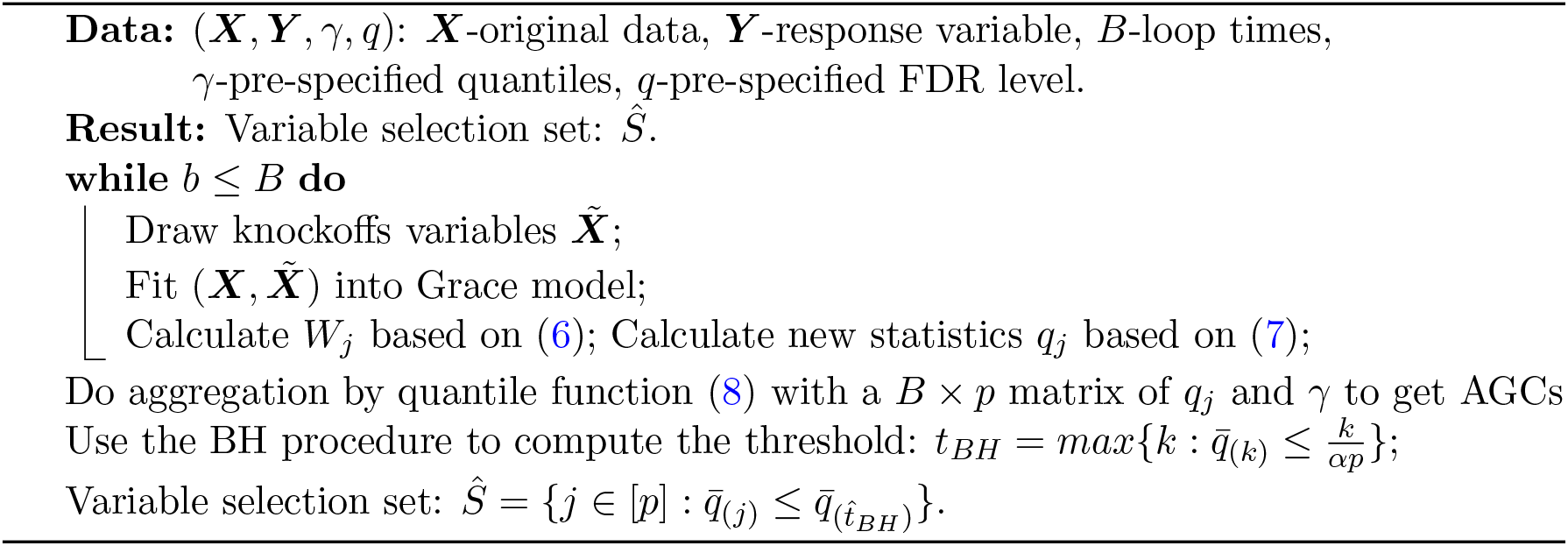

As an illustration in Candes et al. (2018), the primary objective of the knockoff filter procedure was to build an as permissive as possible data-dependent threshold. The threshold can provide a controllable estimation of FDR to ensure model-X knockoffs property. Nguyen et al. (2020) was based on a reformulation of the original knockoff inference and developed an intermediate feature statistics to replace the feature statistics of Candes et al. (2018). Specifically, AKO still provided a data-dependent threshold based on the specification in Candes et al. (2018) which is used by Grace-AKO.

## 3 Simulation Studies

In this section, we performed a wide range of simulation studies to evaluate our proposed method, Grace-AKO, and compared it to the network-constrained regularization procedure (namely Grace) (Li and Li, 2008). We supposed that there were 10 transcription factors (TFs) and each regulated 10 genes. The graphical structure included *g* unconnected regulatory modules with p genes in total and edges linked each of the TFs and 10 genes regulated. Here we denoted the first 44 genes related to the response variable **y**. We assumed that the data were simulated from the following settings:

1. **y = X*β* + *ϵ***, where

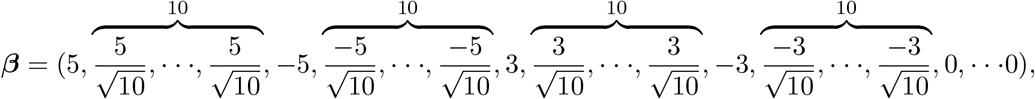

and *ϵ* was generated from *N*(0, *σ*^2^);
2. The noise level was denoted as 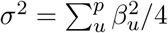;
3. For each expression level TF, *X* was drawn from normal distribution: ***X**_g_* ~ *N*(0, 1), and conditional on the TF, the expression levels of genes which regulated to the specific TF were drawn from a conditional normal distribution with correlation of 0.7.

We set *n* = (100, 200, 300), *g* = (10, 20, 40, 60) and *p* = (110, 220, 440, 660), where *g* represented the total number of unconnected regulatory modules, to simulate 12 settings in total. Figure 1 shows the sub-matrix in order to provide a more accurate depiction of the graphical structure, given that every 11 of the first 44 variables form an identical sub-network. As the empirical research in Nguyen et al. (2020) demonstrated, performance was stable and robust once the iteration *B* approached 25 times. In the meanwhile, FDR could be empirically controlled under the pre-specified level. Therefore, we fixed *B* = 25, and *γ* = 0.1 in all scenarios. Additionally, we set the mFDR control level *α* at 0.1. According to Li and Li (2008), Grace performed better than the Lasso (Tibshirani, 1996) and the elastic net (Zou and Hastie, 2005) in term of the combination of || · ||_1_ and || · ||_2_ and the Laplacian matrix **L**. We thus focused on the comparison of Grace-AKO and Grace in our simulation studies. We assessed the regression performance using the average values of mFDR, standard errors (calculated by Monte Carlo method), and the number of variables selected.

**Figure 1:**
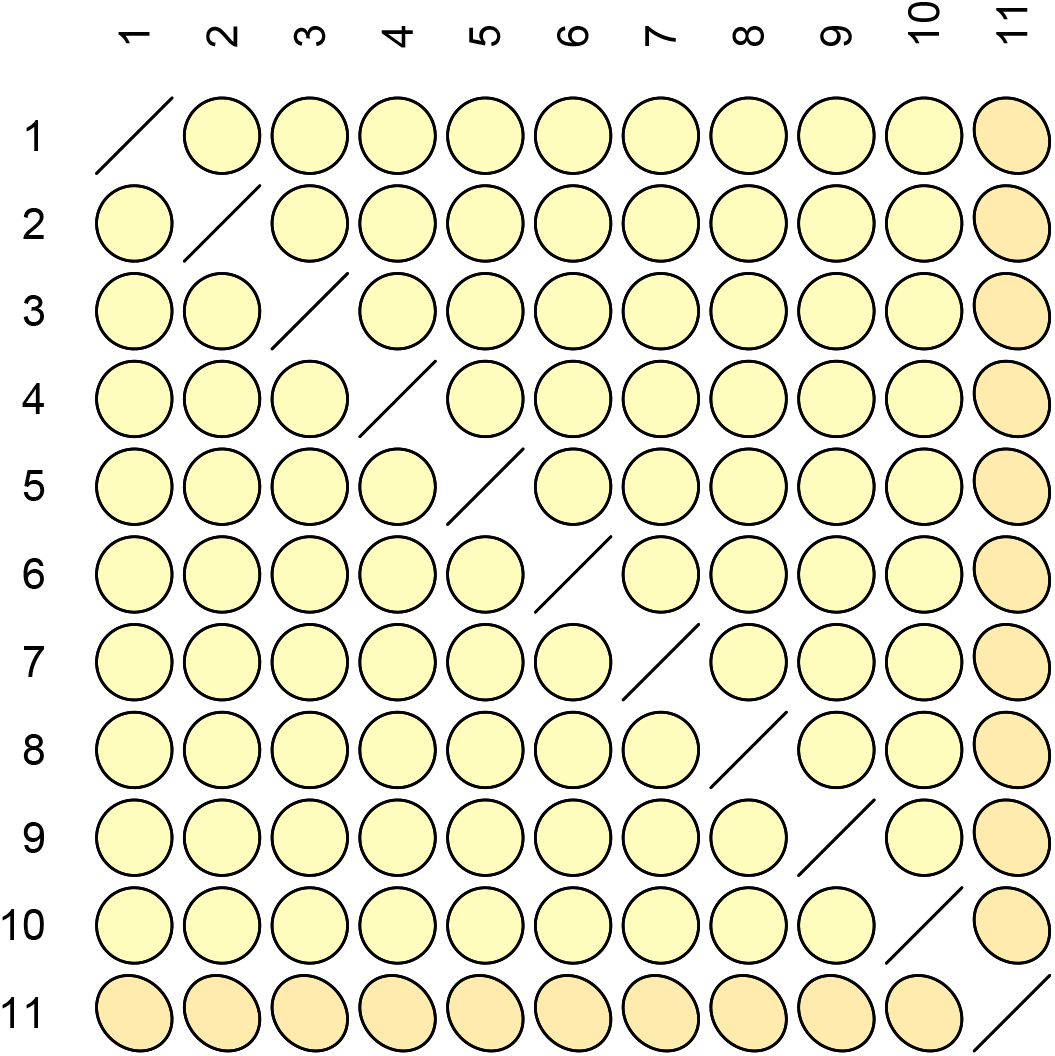
Figure of 11×11 sub-matrix of Laplacian matrix. The upper half of the matrix is color-coded to indicate correlation. The below half of the matrix is numeric-coded to indicate correlation.

To tune the parameters, we applied 10-fold cross-validation. The values of tuning parameter *λ*_L_ were specified from the range of 0.1 to 2.0 with a step size of 0.1, which ensured that the **L** matrix was non-negative. Moreover, the values of *λ*_1_ were drawn from the range of 110 to 200 with a step size of 5. The validation set of *λ*_2_ ranged from 1 to 10 with a step size of 1. Each simulation setting was repeated for 30 times.

As Table 1 showed, the mFDR values of Grace-AKO were controlled at the pre-specified level, and Grace could not control mFDR in finite-sample settings. Furthermore, as the data dimension *p* increased with a fixed sample size *n*, the mFDR for Grace increased which even reached at 0.67. However, Grace-AKO could always control the mFDR under 0.1. Moreover, the standard errors for Grac-AKO were always smaller than Grace’s, which indicated that Grace-AKO could also improve the stability. In Figure 2, we depicted a boxplot for the findings when *n* = 300. The upper figure’s black line represented the pre-specified FDR level. Grace, in particular, had inflated power at cost of high mFDR.

**Figure 2:**
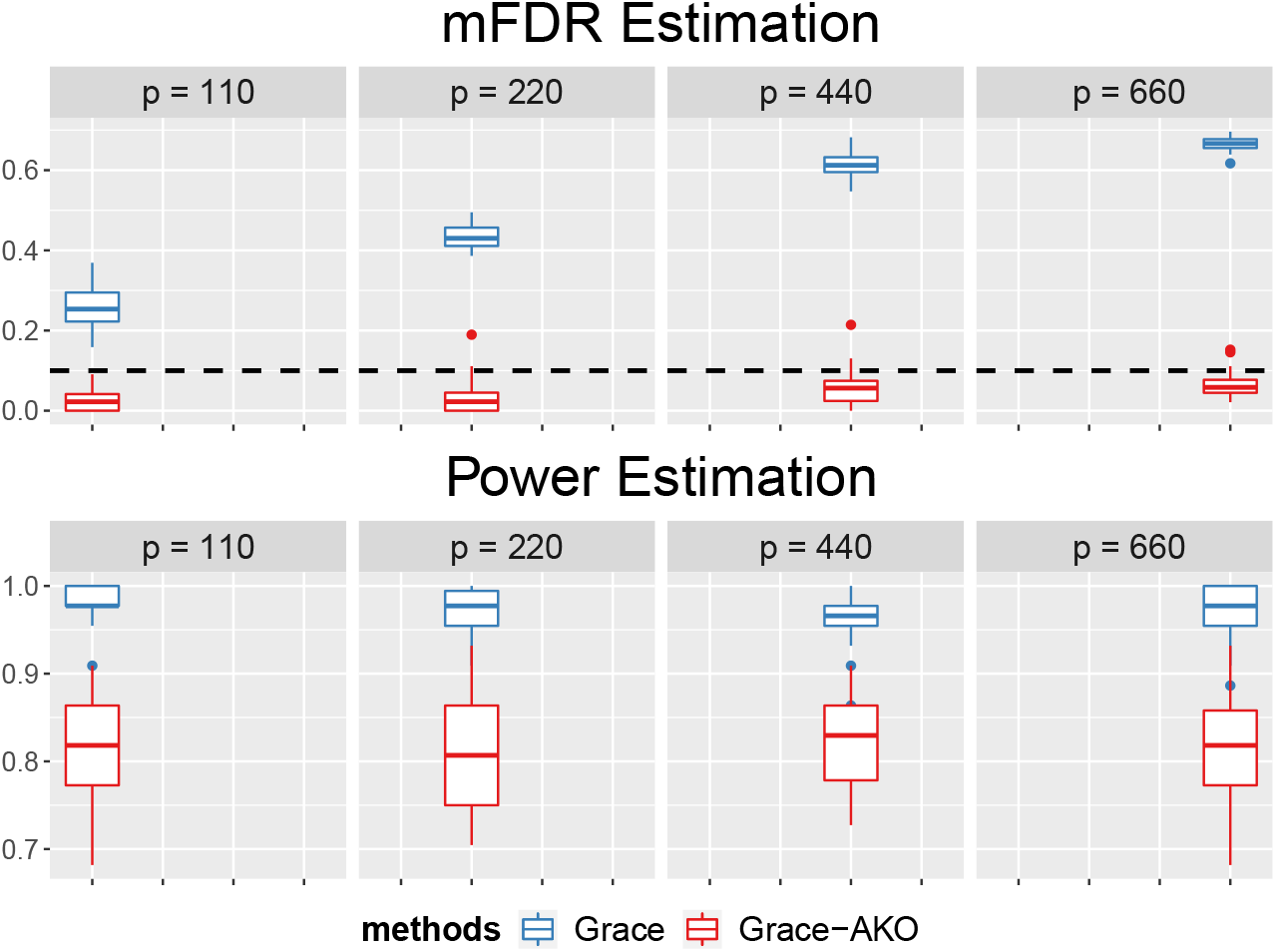
Figures of mFDR and Power for Grace-AKO v.s Grace model when *n* = 300. The upper figure plots mFDR and lower figure plots power. The red color represents our new method, Grace-AKO and the blue color represents Grace. Each setting has been replicated for 30 times. The black dashed line is the pre-specified mFDR level which equals to 0.1.

**Table 1:** The mean and standard errors (in brackets) of modified false discovery rate (mFDR) over 30 replications for Grace-AKO v.s. Grace model with a pre-specified mFDR level at 0.1.

| $p$ | $n = 100$ | | $n = 200$ | | $n = 300$ | |
| --- | --- | --- | --- | --- | --- | --- |
|  | Grace-AKO | Grace | Grace-AKO | Grace | Grace-AKO | Grace |
| 10 | 0.01 (0.018) | 0.15 (0.032) | 0.03 (0.032) | 0.22 (0.049) | 0.03 (0.025) | 0.26 (0.053) |
| 20 | 0.02 (0.027) | 0.26 (0.045) | 0.02 (0.023) | 0.37 (0.045) | 0.03 (0.040) | 0.43 (0.010) |
| 40 | 0.03 (0.030) | 0.24 (0.062) | 0.04 (0.030) | 0.48 (0.040) | 0.06 (0.047) | 0.61 (0.030) |
| 60 | 0.04 (0.026) | 0.40 (0.053) | 0.03 (0.026) | 0.55 (0.032) | 0.06 (0.034) | 0.67 (0.018) |

As shown in Table 2, the number of selected variables varied significantly between these two models. The number of true variables was fixed at 44. When *n* and *p* increased, Grace selected many false discoveries. Notably, when *n* = 300 and *p* = 660, Grace identified 138 candidate variables. This was the reason why Grace’s power was inflated in Figure 2. Grace-AKO showed considerably more stable performance, and it can guarantee finite-sample FDR control under all data settings with slightly conservative power.

**Table 2:** The average numbers of variables selected for Grace-AKO v.s. Grace over 30 replications with a pre-specified mFDR level at 0.1. The number of true variables is 44.

| $p$ | $n = 100$ | | $n = 200$ | | $n = 300$ | |
| --- | --- | --- | --- | --- | --- | --- |
|  | Grace-AKO | Grace | Grace-AKO | Grace | Grace-AKO | Grace |
| 10 | 24 | 45 | 33 | 56 | 37 | 62 |
| 20 | 25 | 52 | 32 | 71 | 37 | 83 |
| 40 | 24 | 59 | 34 | 90 | 39 | 118 |
| 60 | 24 | 63 | 32 | 101 | 39 | 138 |

We further assessed the robustness and single knockoff (model-X knockoffs) performance on simulated data (*n* = 100, *p* = 110, and a pre-specified mFDR = 0.1). To evaluate the robustness of Grace-AKO, we randomly selected 20 vertices from the first 44 elements (true candidate genes) of the Laplacian matrix and set their degrees to zero. Additionally, we set the first and third TFs to have false degrees of 1 and 4, respectively. The mFDR and TPP of Grace-AKO were 0.013 and 0.516, respectively. It indicated that Grace-AKO could still control the mFDR under the pre-specified FDR level (mFDR = 0.1), despite the fact that some information of graphical structure was misspecified for the true variables. Moreover, prior researches demonstrated that the findings might be robust to the misspecification of the graphical structure (Li and Li, 2008; Wei and Li, 2007, 2008). As there were few genes associated with the response variable, the majority of coefficients would be zero. In addition, we examined the performance of Grace incorporating with model-X knockoffs, termed as Grace-KO over 200 simulations. We conducted simulations with 200 replications when *n* = 100 and *p* = 110. The results were reported in Table **??**. We observed that Grace-KO and Grace-AKO were both able to control the mFDR under the pre-specified mFDR level, and Grace-AKO shown more stable performance with a lower standard deviation. Moreover, Grace-KO performed more conservatively than Grace-AKO, which was able to identify fewer candidate genes. Furthermore, we also assessed the computational cost of Grace-AKO when *n* = 100 and *p* = 110. The knockoff generation and inference steps of Grace-AKO could be conducted by parallel running, which took about 30 seconds with a server of Intel Xeon Silver 4116 CPU 2.10 GHz and 64 GB RAM memory. Consequently, we concluded that Grace-AKO was robust to the incorrect information of the graphical structure and was more powerful in identifying candidate variables.

## 4 Application to the TCGA Prostate Cancer Data

To demonstrate the usefulness of Grace-AKO, we applied it to a gene-expression data of prostate-specific antigen (PSA) level from The Cancer Genome Atlas (TCGA) program. The TCGA program is a landmark cancer genomics program with over 11,000 cases of primary cancer samples spanning 33 cancer types (Koboldt et al., 2012). Additionally, the Kyoto Encyclopedia of Genes and Genomes (KEGG), as a public database, contains rich information about the graphical structures of genes (Kanehisa and Goto, 2000). It contributes to understanding various aspects of biological systems and pathways. Prostate cancer is the most common malignancy in mid-aged males and the second leading malignancy (Stangelberger et al., 2008). This external graphical structure is represented by a penalty weight matrix, which is the Laplacian matrix ***L*** constructed in equation (3). Wang et al. (2018) indicated that metastatic prostate cancer remained incurable even in patients who finished intensive multimodal therapy. It is an urgent challenge to propose a novel approach for disease management via identifying prognostic determinant genes. Moreover, Reiner et al. (2003) indicated that the statistical significance of differential expression might require abundant experiments, and the probability of type I error increased as the multiple hypotheses were tested. In this article, we thus were more concerned with the accuracy of gene selection in tumor investigations (assessed by mFDR) than the power of detection.

We first removed the samples with missing measurements and then encoded PSA pathways (Kim et al., 2018) from the KEGG to construct the normalized Laplacian matrix for Grace and Grace-AKO. We finally obtained the data with sample size *n* = 339 and dimension *p* = 5, 947. We denoted the PSA level as our response. To reduce computational demands, we first performed variable screening via correlation learning following Fan and Lv (2008). The final sample size and dimension were *n* = 339 and *p* = 600, respectively, consisting of the data structure in simulation studies. We then standardized the explanatory variables and centered the response. The parameter *λ*_1_ ranged from 1 to 40 length out as 4, and *λ*_2_ was among 1, 2, 3, 4, and 5. We denoted the target FDR level at 0.2 and *B* = 25 for the iterations of AKO procedure. The quantile point was *γ* = 0.3. For a fair comparison, we both conducted 10-fold cross-validation to select the tuning parameters. We showed the details in Figure 3.

**Figure 3:**
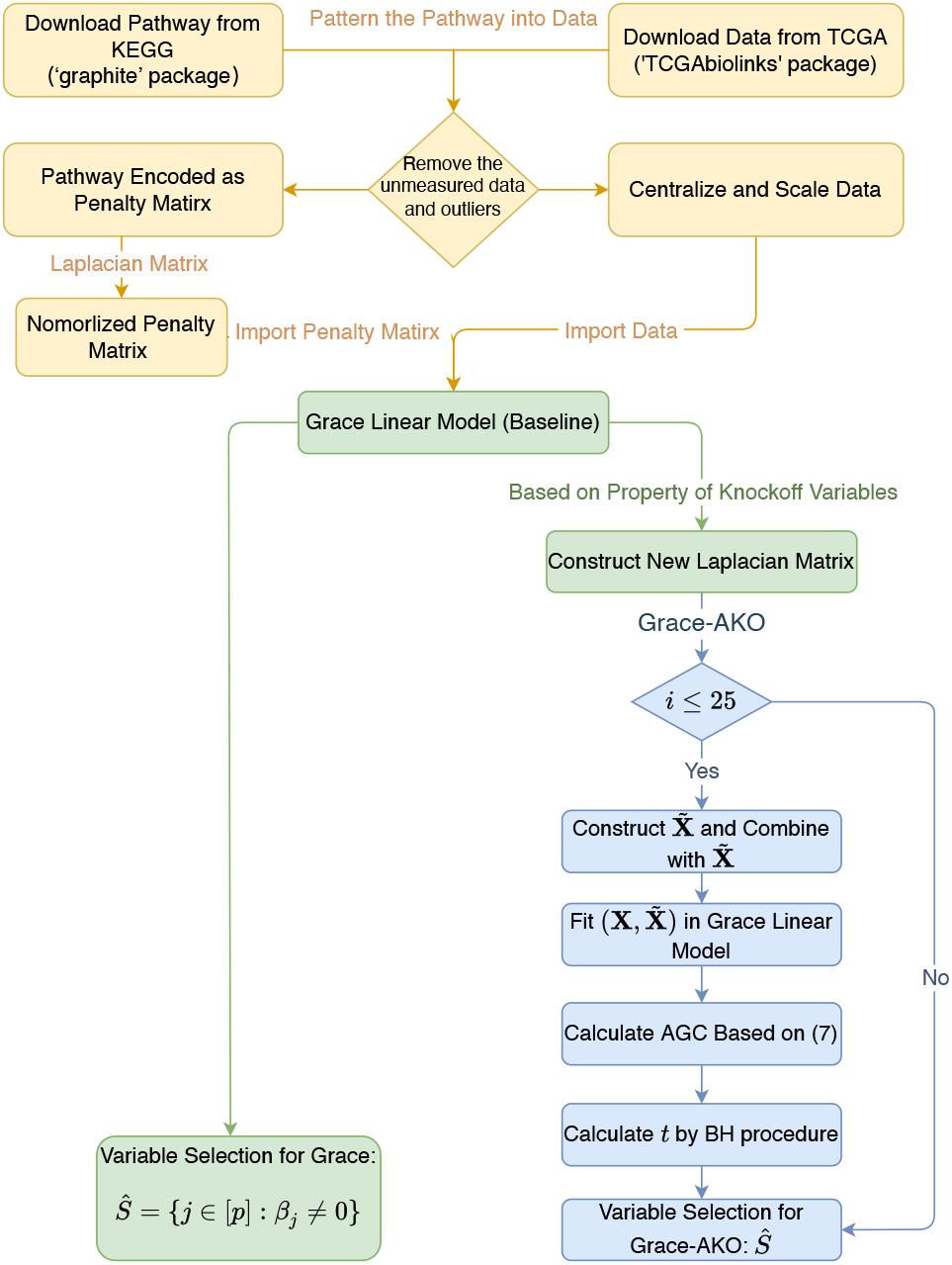
Flowchart of Grace-AKO. These two procedures are differentiated by colors, in which the green one is Grace and the blue one is Grace-AKO. First, we encode the Laplacian matrix account for the network structure and sample model-X knockoffs. Second, we compute the feature statistics LCDs. Third, we aggregate the LCDs to get AGCs. Fourth, we select the variables whose AGCs satisfy the requirements.

There were 90 genes selected by Grace in total, and 47 genes selected by Grace-AKO. Li and Li (2010) indicated that Grace might lose some accuracy when the coefficients’ signs are different. After checking the intersection of variable selection sets, we found that the genes selected by Grace-AKO were a subset of Grace’s. However, due to the inflated mFDR in the simulation studies, the results of Grace might identify some false discoveries. Moreover, previous studies confirmed 35 out of 47 candidate genes detected by Grace-AKO were related to prostate cancer. Some details of them were listed in Table 3. For example, IL9 was recently assigned as an essential gene in tumor immunity, and AK6 was already regarded as a biomarker in prostate cancer treatments (Liu et al., 2015; Wan et al., 2020). HLA-DRB5 was highly related to MHC-II genes, which were the related genes of prostate cancer, and so was the CHRNB2 (Axelrod et al., 2019). Moreover, a high concentration of IFNA2 was related to advanced prostate carcinoma (Erb et al., 2013). PLK1 was over-expressed in many cancers, including prostate cancer, and scientists found that translation of the PP2A-PLK1 SDL interaction caused the expression of PLK1 and PP2A, which were commonly regarded as a biomarker in the cancer cells (Cunningham et al., 2016). Melloy (2019) indicated the percentage of the case with alteration of ANAPC7 achieved beyond 5.21 percentage points. The results demonstrated that the APC/C had a profound effect on cancer survival. MYC in 2010 had already been confirmed to be affected by the loss of the tumor in Koh et al. (2010).

**Table 3:**
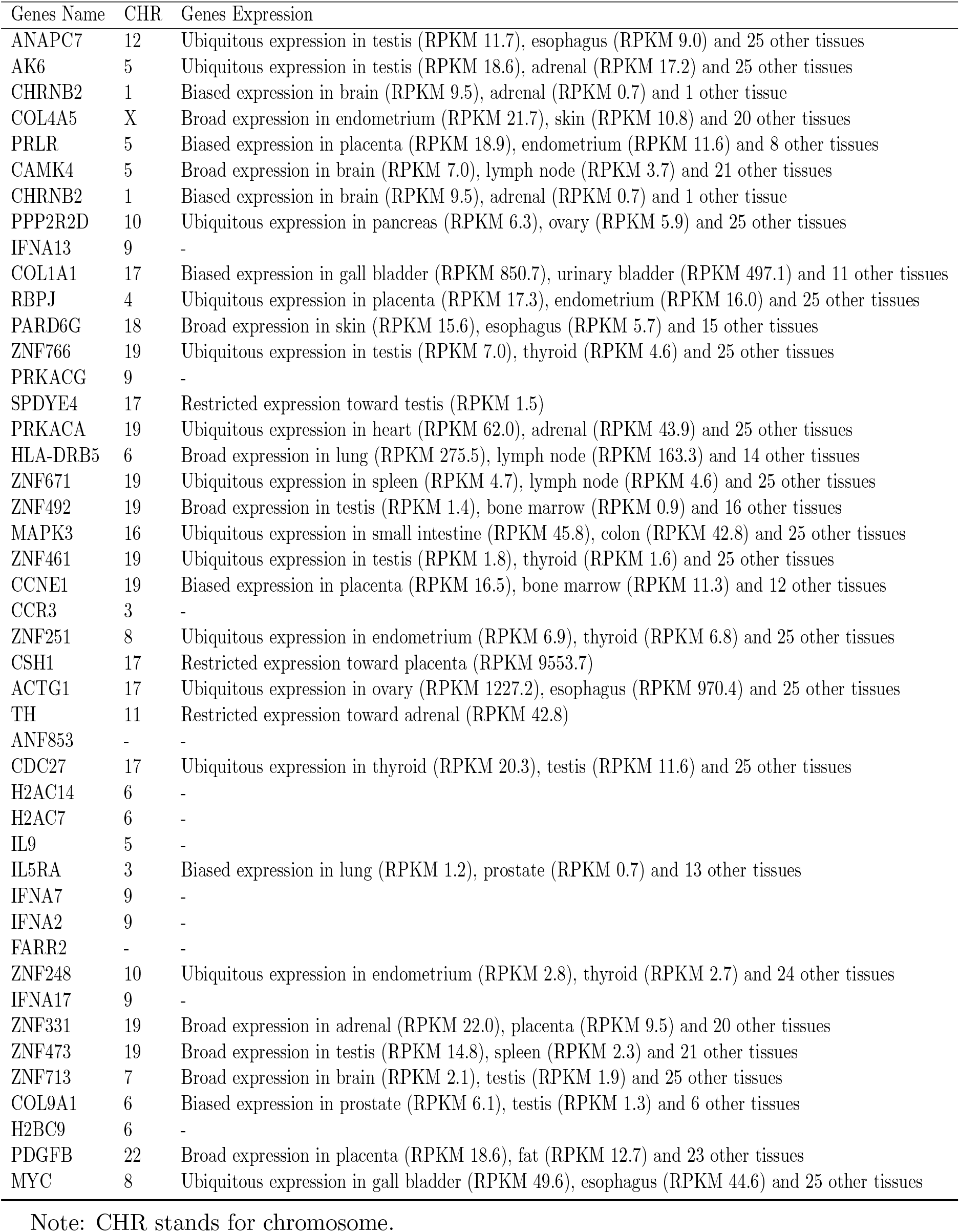
Genes Selected by Grace-AKO in the application of prostate cancer and PSA pathways.

Furthermore, there were also some subnetworks in our findings. H2BC9, H2AC14, and H2AC7 consisted of a small subnetwork. Moreover, IFNA7, IL9, IL5RA, IFNA2, PDGFB, and KIT comprised a subnetwork of validated pathways. Except for KIT, the remaining genes were investigated as possible prostate cancer therapy genes. Additionally, there was a pathway between CAMK4 and PRKACF, where CAMKK2 was a significant androgen receptor target for prostate cancer tumor growth, according to Dadwal et al. (2018). MYC was linked with RXRA. In Ray et al. (2020), RXRA, which was discovered as a novel target of miR-191, was conserved in a cell line derived from radio recurrent prostate cancer.

## 5 Conclusions

This article introduces Grace-AKO to perform variable selection by incorporating the network-constrained penalty (Li and Li, 2008) and AKO (Nguyen et al., 2020). In contrast to the conventional variable selection process, Grace-AKO applies a normalized Laplacian matrix to encode the graphical structures between the potential genes or the gene products. It applies the *L_1_* penalty to selected variables and the *L_2_* penalty to degree-scaled differences of coefficients concerning the graphical structure. Moreover, our proposed Grace-AKO guarantees of FDR control in finite-sample settings by identifying variable employing multiple knockoff vari-ables. Grace-AKO addresses the instability of model-X knockoffs by incorporating a statistical aggregation procedure and introducing a new feature statistics AGC. The simulation results in-dicated that Grace-AKO had superior performance in finite-sample FDR control in a wide range of simulation settings. In order to control the finite-sample FDR, Grace-AKO would be slightly conservative in terms of power (Gimenez and Zou, 2019). Furthermore, the proposed general framework for variable selection with finite-sample FDR control can be broadly extended to other existing penalties (e.g., the Lasso penalty (Tibshirani, 1996) and the elastic net penalty (Zou and Hastie, 2005)).

## Notes

### Competing Interest Statement

The authors have declared no competing interest.

### Summary of Updates

Added some details about the proposed method and new simulations

